# Determining Temporal Stability in Dominance Hierarchies

**DOI:** 10.1101/692384

**Authors:** C. Vilette, T.R. Bonnell, S.P. Henzi, L Barrett

## Abstract

The importance of social hierarchies has led to the development of many techniques for inferring social ranks, leaving researchers with an overwhelming array of options to choose from. Many of our research questions involve longitudinal analyses, so we were interested in a method that would provide reliable ranks across time. But how does one determine which method performs best?

We attempt to answer this question by using a training-testing procedure to compare 13 different approaches for calculating dominance hierarchies (seven methods, plus 6 analytical variants of these). We assess each method’s performance, its efficiency, and the extent to which the calculated ranks obtained from the training dataset accurately predict the outcome of observed aggression in the testing dataset.

We found that all methods tested performed well, despite some differences in inferred rank order. With respect to the need for a “burn-in” period to enable reliable ranks to be calculated, again, all methods were efficient and able to infer reliable ranks from the very start of the study period (i.e., with little to no burn-in period). Using a common 6-month burn-in period to aid comparison, we found that all methods could predict aggressive outcomes accurately for the subsequent 10 months. Beyond this 10-month threshold, accuracy in prediction decreased as the testing dataset increased in length. The decay was rather shallow, however, indicating overall rank stability during this period.

In general, a training-testing approach allows researchers to determine the most appropriate method for their dataset, given sampling effort, the frequency of agonistic interactions, the steepness of the hierarchy, and the nature of the research question being asked. Put simply, we did not find a single best method, but our approach offers researchers a valuable tool for identifying the method that will work best for them.

**Highlights:**

- All ranking methods tested performed well at predicting future aggressive outcomes, despite some differences in inferred rank order.
- All ranking methods appear to be efficient in inferring reliable ranks from the very start (i.e., with little to no burn-in period), but all showed improvement as the burn-in period increased.
- Using a common 6-month burn-in period, we found that all methods could predict aggressive outcomes accurately for the subsequent 10 months. Beyond this threshold, accuracy in prediction decreased as the testing dataset increased in length.
- Switching to a data-driven approach to assign k-values, via the training/validation/testing procedure, resulted in a marked improvement in performance in the modified Elo-rating method.

## INTRODUCTION

“Which of these methods is the best?” was the question that set us on the path of this paper. The lack of a clear answer convinced us to write it. We were conducting an analysis of longitudinal data from our long-term study of vervet monkeys and wanted a method of calculating dominance ranks that would give good reliability; that is, we wanted to know whether the hierarchy inferred today could reliably predict a later date. If you are looking to calculate dominance ranks, however, you will soon have “*l’embarras du choix*”: there is such an such an overwhelming array of options that you may not even know where to start, let alone decide which one to choose. Our analyses revealed that we couldn’t hope to provide a straightforward answer—there is no absolute best method—but we did develop an approach that enables researchers to find the method that will provide the best fit for their data. Specifically, we demonstrate the use of a training-testing procedure that not only enables the most appropriate method to be identified, but also offers greater insight into the structure and dynamics of a given dataset. In what follows, we compare 13 different approaches for calculating dominance hierarchies (seven methods, plus six analytical variants of these). We assess the methods’ performance, their efficiency, and the extent to which the calculated ranks obtained from the training dataset reliably predict the outcome of observed aggression in the testing data set.

### A Brief Survey of Analytical Methods for calculating Dominance Ranks

The analytical importance of hierarchies to our understanding of social organisation has driven the development of several techniques for inferring dominance hierarchies from observational data (de Vries 1998; reviewed in Bayly, Evans, & Taylor 2006; Whitehead 2008; Briffa et al. 2013), as well as extensive research directed at their formation and maintenance (e.g. Parker 1974; Dugatkin & Earley 2004; Sasaki et al. 2016). At present, there are seven methods that either have been commonly used to calculate dominance hierarchies, or newly offered as improvements to existing methods.

Until recently, all ranking methods were based on interaction matrices. Using this approach, it is possible to distinguish two ways of quantifying dominance relationships. In the first, the dominance matrix is reorganized such that some numerical criterion is minimized or maximized (e.g., Inconsistencies and Strength of Inconsistencies method (I&SI): de Vries 1998; ADAGIO (‘approach for dominance assessment in gregarious species’): Douglas et al. 2017). The second approach aims to provide a suitable measure of individual overall success, from which a rank order can be derived directly (e.g. David’s score: David 1987, Gammell et al. 2003; Elo-rating: Elo 1978, Neumann et al. 2011). The I&SI (method 1) and the David’s score (method are the two most commonly used ranking techniques. Despite their usefulness, they share a drawback common to all matrix-based methods. Namely, they gather all observed interactions within a particular time period to determine a single quasi-static ranking, with the result that the obtained ratings depend directly on the number of individuals present in the matrix itself. Thus, any observed fluctuation in rank across two time periods may simply be a consequence of demographic events rather than variation in competitive abilities. Hence, these methods fail to provide information about how ranks may change or are maintained over time.

In order to address actual, rather than “demographic”, changes in a rank hierarchy, a common approach has been to compare hierarchies across different periods, or before and after specific events, such as the integration of a new individual (Arseneau-Robar et al. 2017). However, this still fails to track any potential continuous variation in the rank order of subjects (c.f. Neumann et al. 2011; Newton-Fisher 2017). At the same time, breaking down the assessment of rank hierarchies into shorter time windows is itself problematic, as shorter sampling periods may result in a lack of data from which to infer a reliable hierarchy (Goffe, Fischer & Sennhenn-Reulen 2018). The I&SI method has an additional downside in that it seeks to produce a linear rank order, regardless of whether the society in question is expected to show a linear hierarchy, or whether the data fit the assumption of linearity. These methodological flaws have led to the emergence of two alternative approaches.

First, given that simple linearity is not likely to characterise the dominance structures of many animal societies, Fujiii et al. (2015) developed a network-based method, called Percolation and Conductance (P&C). This permits nonlinear structure to emerge via estimates of network directional consistency in the flow of dominance interactions, and the detection of blocks of dominance ambiguity that are indicative of nonlinear segments of a hierarchy. This technique uses paths within the agonistic network to generate an individual’s probability of winning against all other individuals. Despite overcoming the linearity assumption, P&C nevertheless follows the two previous methods in being reliant on interaction matrices. This brings us to the second alternative, which enables a more dynamic assessment of rank over time.

This method can be traced to Neumann et al. (2011), who argued that matrix-based methods for rank assessment often cannot be applied to highly dynamic animal societies, or to sparse datasets. With this in mind, he introduced the Elo-rating method (method 4), derived from its application in competitive games, and famous for its use in rating chess-players (Elo 1978). This method enables the rating process to continue despite changes in group-composition, thus circumventing one of the other drawbacks of matrix-based methods (viz. I&SI, DS and P&C). In other words, Elo-ratings are based on the sequence in which interactions occur, with ranks continuously updated. This has the advantage of allowing ratings to span changes in group-composition.

Despite these improvements, Elo-rating also has its limitations. One fundamental problem is that, in the absence of any knowledge of prior dominance relationships, the method assigns all individuals the same initial Elo-rating score, which is then updated as interactions are added across the observation period. Consequently, a “burn-in” period is necessary so that sufficient observations can accumulate and enable the modeled rankings to catch up with the computation of reliable ratings of dominance relationships (Albers and de Vries 2001; Neumann et al. 2011). Both Albers & de Vries (2001) and Neumann et al. (2011) are vague about how long this process might take, probably because the duration of the burn-in will vary with the frequency of agonistic interactions (Newton-Fisher 2017).

Another issue with Elo-rating is the assumption that all agonistic interactions entered into the model are equivalent in their potential influence on rank trajectories. In rating subjects, the variable k is used to determine the degree to which each interaction influences the future rank trajectory of both winner and loser. In other words, it determines the number of rating points that an individual gains or loses after a single encounter (Neumann et al. 2011). Newton-Fisher (2017) argues that holding k constant makes the implicit assumption that, as long as a clear winner and loser can be identified, variation in the intensity of aggression does not influence social dominance rank or rank trajectories.

As Elo-rating has received increasing interest, there have been a number of recent attempts to address the difficulties associated with its use (e.g. Foerster et al. 2016a; Newton-Fisher 2017; Goffe, Fischer & Sennhenn-Reulen 2018; Sánchez-Tójar, Schroeder & Farine 2018). This has resulted in the development of a method to calculate dominance, referred to as the modified Elo-rating method (method 5). Specifically, Newton-Fisher (2017) presents two developments of Neumann et al.’s (2011) R function to improve its efficiency: (i) the incorporation of prior history and (ii) the recognition of differing intensities of aggression in agonistic interactions. Using data from male chimpanzees (*Pan troglodytes*), Newton-Fisher (2017) showed that incorporating even limited prior knowledge of dominance ranks substantially reduced the required burn-in period and improved the effectiveness of Elo-rating in resolving hierarchical structure. In addition, allowing k to vary further improved the Elo-rating models. Taking another approach to the limitations of the burn-in period, Goffe, Fischer & Sennhenn-Reulen (2018) used “partial pooling”, which rests on the assumption that all initial ratings are sampled from the same distribution with a shared variation parameter σ. This Bayesian Inference (BI) approach (method 6) facilitates the estimation of initial ratings, as well as the value of k.

Finally, one of the aims of the original Elo-rating was to track dynamic changes in rank. However, most behavioural studies assume that individual dominance rank is relatively stable over time (e.g. Poisbleau, Guillon & Fritz 2010). With this in mind, Sánchez-Tójar, Schroeder & Farine (2018) suggested a modification to the original Elo-rating based on randomizing the order in which interactions occurred: the randomized Elo-rating method (method 7). They showed that randomizing the order in which interactions occurred (1000 times), and estimating mean individual ranks, improved performance compared to the original Elo-rating, particularly when hierarchies were not unduly steep. This finding supports the idea that randomizing interaction order has a beneficial effect when species’ social dynamics are relatively stable and rank acquisition is achieved by the pattern of maternal rank inheritance. The principles underlying all seven methods are outlined in Figure 1.

**Figure 1:**
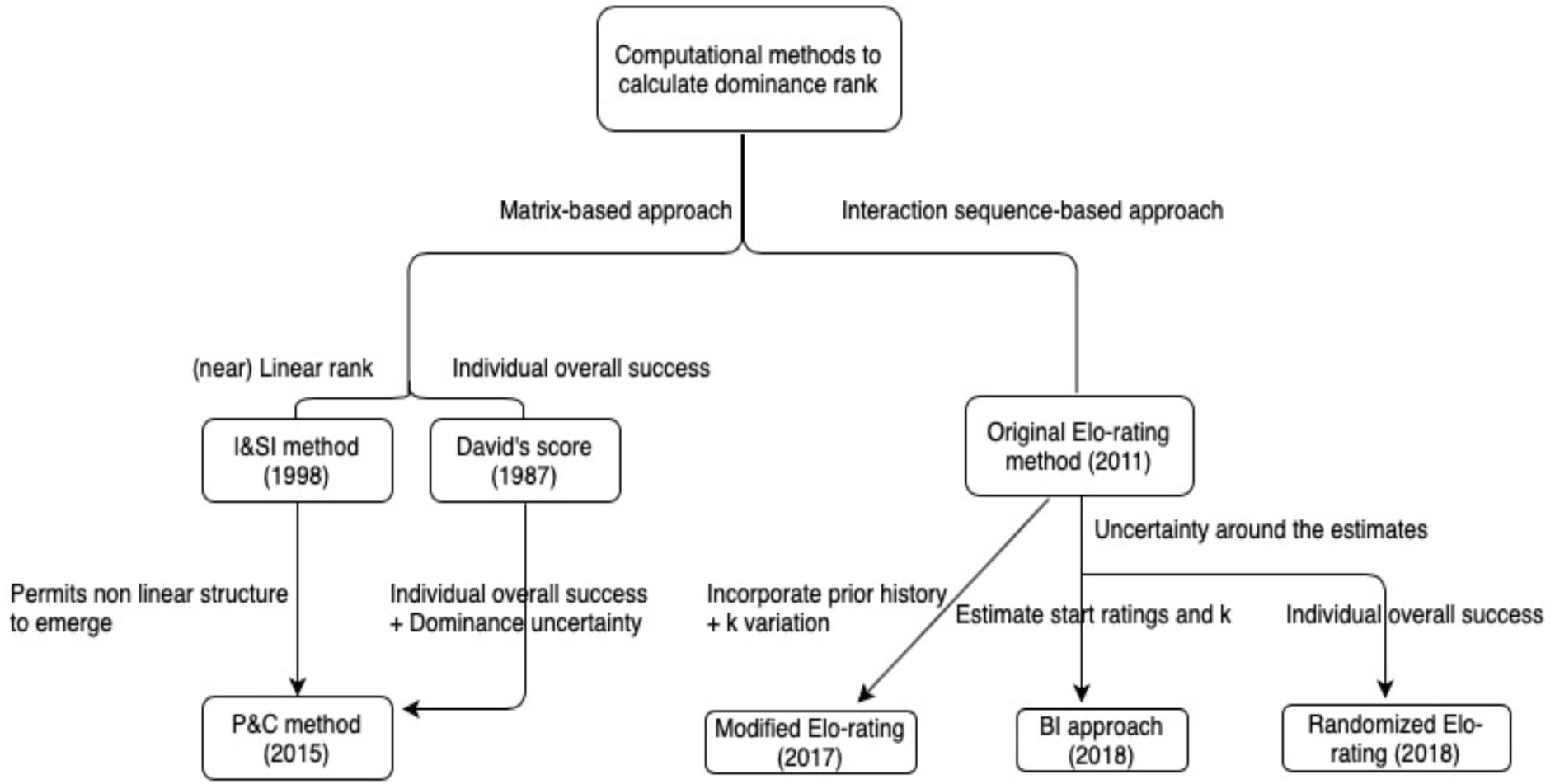
Main-guiding principles of each cited methods.

Several attempts have been made to assist researchers in selecting the most appropriate method for quantifying dominance ranks, with most focusing on comparing the level of agreement among several methods when applied to real datasets (e.g. Balasubramaniam et al. 2013; de Vries 1998; Gammell et al. 2003; Neumann et al. 2011). Sánchez-Tójar, Schroeder & Farine (2018) argue that, while cases where two or more methods closely match one-another could signify that they are robust, it might also mean that they suffer from a common bias and, if so, such comparisons can provide no information about their accuracy. Consequently, they recommend simulating artificial datasets containing individuals of known rank, as well as simulating interactions among those individuals under different scenarios of known steepness, and then testing the validity of the method(s) by correlating the inferred hierarchy to the known hierarchy.

Here we take an alternative, but complementary, approach. Using agonistic data from a long-term study of vervet monkeys (*Chlorocebus pygerythrus*), we generate training-testing datasets in order to determine the efficiency of the seven listed methods (along with several variants of particular methods). The notion that dominance hierarchies reduce uncertainty about the outcomes of contests between group members (Beaulieu et al., 2014; Mendonça-Furtado et al., 2014) assumes that the state of the hierarchy at a given time will be predictive of future interactions (Strauss & Holekamp, 2019). We test this assumption by assessing whether individual ranks, obtained from the training dataset, can predict the future aggressive outcomes that occur in the testing dataset. We proceed as follows.

First, we explore each method’s performance by determining the proportion of accurate predictions in aggressive outcomes across the testing dataset. That is, we assess whether the ranks obtained from a large training dataset are in agreement with the outcomes of aggressive interactions present in our testing dataset. Given that, over time, groups undergo changes in their social dynamics due to changes in the social and ecological environment, both of which may underpin shifts in rank, we predict that interaction sequence-based approaches will perform better in deducing ranks than the matrix-based ones, as the former methods take into account the temporal dynamics of the social hierarchy (Neumann et al. 2011; Williamson, Lee & Curley 2016; Newton-Fisher 2017; Goffe, Fischer & Sennhenn-Reulen 2018).

Following this investigation, we assess each method’s efficiency in terms of the amount of data needed to obtain accurate estimations. A burn-in period is necessary for sufficient observations to accumulate and enable a stable rank position to be calculated. However, an excessive amount of data may not give a representative picture of the current state of play within a group, because old, and potentially out-of-date, information will be included in the calculations. In order to find the right trade-off, we determine the necessary duration of the burn-in phase for each method by repeatedly reducing the length of training dataset size by 2-month increments. To allow for comparison, we kept the testing dataset constant. To put this in concrete terms, we maintain the same end date for the training dataset, while varying its start date. Thus, as the training dataset decreases in size, only the most recent observations are included. At each reduction in length, individual ranks are calculated and tested against the testing dataset. We expect to see the reliability of predicted ranks to increase with the amount of data used until a threshold is reached (representing the accuracy limits of older information being included in the calculation), following which ranks should decrease in their reliability.

Finally, we analyze how well future aggressive outcomes can be predicted by values calculated during an earlier period. To do this, we reversed the above process, keeping the training dataset constant, while sequentially extending the testing dataset size by 2-month increments, until we achieved a 30-month testing dataset. Our interest here was to assess the length of time it would be possible to use previously calculated ranks with minimal loss of accuracy. As mentioned above, rank positions within a hierarchy can change over time due to temporal variation in demographic and ecological conditions. Thus, we expect that aggressive outcomes will be predicted with high accuracy initially and will then manifest a constant decay as the accuracy of ranks decreases through time.

After investigating these seven methods, we conclude our study by proposing our own modification to the modified Elo-rating that helps to improve its performance.

## METHODS

### a. Study site and subjects

Data used for these analyses were collected between January 2015 and December 2017 as part of a long-term field project at the Samara Private Game Reserve, South Africa (32o22’S, 24o52’E). We used data from one of our three study groups (RBM). All animals were fully habituated and individually recognisable. The study group occupied semi-arid riverine woodland (Pasternak et al. 2013). Group composition varied across the study period (Males: 20-6, Females: 13-8; Juveniles: 33-9; Infants: 11-2).

### b. Behavioural data collection

Agonistic behaviours, identities of participants and interaction outcomes were recorded ad libitum on all group members (i.e., across all sex and age categories). In order to make use of the most diverse and complete dataset, we included agonistic encounters with juveniles and infants and also those that involved coalitions (i.e., where one or more animal comes to the aid of another against a common opponent). Unknown outcomes were discarded. Agonistic behaviours included displacements, threats, chases and bites. The visibility of the habitat, together with the modal presence of more than one observer (McFarland et al. 2014, Henzi at al. 2013), means it is unlikely that there was any systematic bias in the recording of agonism. We recorded 11 323 agonistic interactions between 66 individuals across the 36-month period. The initial training dataset comprised 8 292 interactions, with the testing dataset accounting for the remaining 3 031 interactions.

### c. Methods used to infer ranks

We assessed the performance of three commonly used methods for inferring longitudinal hierarchies: the I&SI method, David’s scores (DS) and the Elo-rating method. We also evaluated the performance of P&C, as well as the three methods derived from Elo-ratings: the BI approach, the randomized Elo-rating and the modified Elo-rating approach (see Table 1 for a summary of the methods) to give a total of seven methods. As noted above, several different statistical packages and options are available for the David’s score and P&C methods, thus we assessed 13 methods in all.

**Table 1:** Summary of the different methods tested in this study

| Method | Approach | Outcome | Analytical details |
| --- | --- | --- | --- |
| I&SI | Reorder interaction matrix iteratively to minimize (1) number of inconsistencies (I) & (2) strength of inconsistencies (SI) | Ordinal rank order | Package: compete<br>Function: isi13<br>nTries= 449 |
| David's score | Matrix based approach in which the relative strength of opponents is taken into account | Individual overall success determined by weighting each dyadic success measure by the un-weighted estimate of the opponent's overall success | Packages: compete, steepness & EloRating<br>Indices: Pij and Dij |
| Percolation & Conductance | Combines information from direct win/loss interactions **and** from indirect pathways to create a matrix of probabilities. The network transitivity determines how much to weight the indirect 'wins' from these pathways | Average dominance probability + dominance uncertainty | Package: Perc<br>MaxLength= 2 and 4 |
| Original Elo-rating | Scores are updated after each interaction depending on the outcome. Individuals start with predefined score while k is a winning/losing shift coefficient that is kept constant | Individual overall success | Package: EloRating<br>Initial scores: 1 000<br>K=100 |
| Modified Elo-rating | Attribute a k value to each of the aggression categories | Individual overall success | Code from<br>Newton-Fisher 2017<br>4 categories of aggression intensity. |
|  |  |  | Lowest starting at k= 200, with k increasing by 25 per aggression intensity |
| Bayesian Inference | Estimates both the most probable rank order + the posterior probability of that order | Start ratings, the Elo-rating winning/losing shift coefficient (k) before computing the scores + the uncertainty around the estimates | Code from Goffe, Fischer & Sennhenn-Reulen 2018 |
| Randomized Elo-rating | Randomized interaction sequence | Individual overall success + measure of uncertainty around the estimates | Package: aniDom<br>n.rand= 1 000 |

All our calculations were implemented in R.

The choices made regarding the analyses of each method are detailed in the Appendix.

### d. Construction of training and testing datasets and comparison of methods

*i. Determining the method’s performance*

We divided our dataset, using the first 80% (2.1.2015 – 25.4. 2017) to train the methods, with testing undertaken on the remaining 20% (26.4.2017 – 31.12. 2017)(Figure 2). This ensured that we always had a training dataset with sufficient observations to infer reliable ranks. This approach is taken from machine learning where it is a common strategy to split data into training and evaluation subsets, usually with a ratio of 70-80 percent for training and 20-30 percent for evaluation. The two datasets are thus distinct as we are only interested in predicting future outcomes and not interpolating ranks. For each method, we calculated dominance hierarchies from our training data.

**Figure 2:**
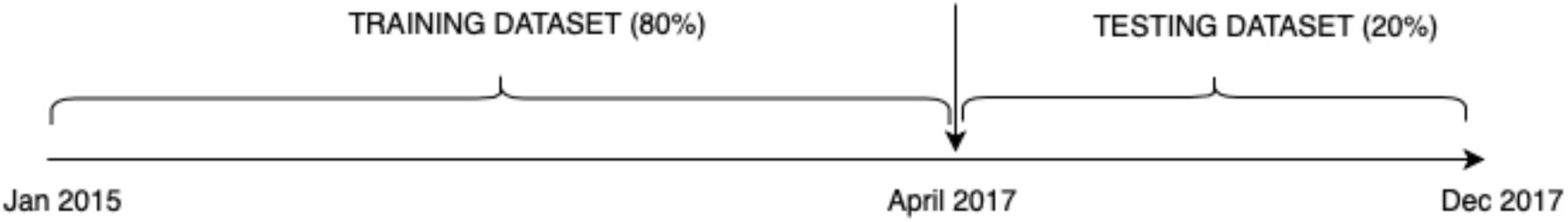
Original approach used to assess method’s performance

It is important to note that, in our case, each model (i.e., ranking method) is fit to the data while a withheld dataset is used to assess the models. As we are interested in the reliability of rank estimates across time, this approach proves to be useful with time series data as it lets us make estimates of how well each model performs in forecasting (i.e., predicting future interaction outcomes). Here, we make the assumption that the process model is the same between the two time periods (i.e., training and testing) and we make “small world” predictions about which method is best for our particular dataset.

We first chose to use visualisation to examine how the rank order inferred by each method differs. Given that the percentage of accurately predicted outcomes results from the inferred rank order, knowing the extent to which rank order varies can help to better understand each method’s performance. To do so, we implemented a hierarchical clustering approach that assembled our results according to their similarity. Initially, each method was assigned to its own cluster. The algorithm then proceeds iteratively, at each stage joining the two most similar clusters, and continuing until there is just a single cluster. In this way, methods that are most similar to each other are combined into branches that are themselves fused higher up in the clustering process. Euclidean distance is used to measure the dissimilarity between each pair of methods. As the estimated ranks were measured on different scales, we standardized the data and used the hclust R function (stats package) to generate this hierarchical clustering.

The obtained ranks for each method were then converted into ordinal ranks to enable comparison across methods. The ordering of these ranks was used to assess how well they matched the outcome of aggressive interactions for each interaction in the testing dataset (yes =1 or no=0). The proportion of accurately predicted outcomes was then translated into a percentage allowing us to assess which methods (if any) outperformed others. We expected more accurate predictions using the more dynamic approaches given changes in the social dynamics of our study group. We excluded from analysis animals that were only present during the testing phase of the dataset, but retained those individuals present only in the training dataset, as they were able to provide information about their opponents.

*ii. Determining the optimal burn-in period*

We assessed the methods’ efficiency by estimating the necessary burn-in period. As noted above, we kept our testing dataset constant while modifying the length of the training dataset. Our original training dataset comprised 28 months, which we reduced sequentially by two months, until only two months were left. To do so, we truncated the dataset starting from January 2015 towards April 2017 (Figure 3).

**Figure 3:**
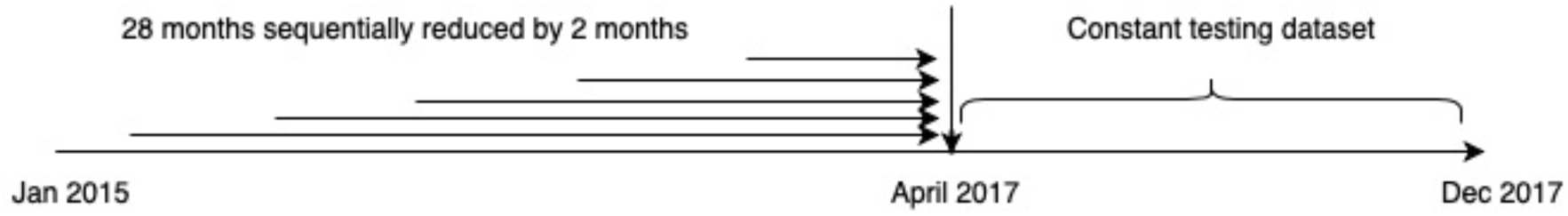
Modified approach used to assess the burn-in period by varying the training dataset length.

At each reduction in size, we computed ranks and assessed these against the testing dataset. The same procedure outlined above was used to calculate the percentage of accurately predicted outcomes. These percentages were then plotted in order to determine the amount of data needed to predict reliable and stable ranks.

*iii. Determining rank stability and predictability across time*

Here, our aim was to determine the length of time over which obtained ranks would continue to accurately predict aggressive outcomes. To do so, we gradually increased our testing dataset size and looked at its impact on the percentage of accurately predicted outcomes. Based on the burn-in period results (see below), we calculated the average optimal training dataset length across all methods. Using the average in this way allowed us to keep the training dataset constant, easing comparisons between the different methods. We then used the remaining data as our testing dataset, and systematically varied its length. We began with the 2-month period that followed on directly from the training phase (July-September 2015) and then sequentially increased the testing dataset by 2 months until the 30-months limit was reached in December 2017 (Figure 4).

**Figure 4:**
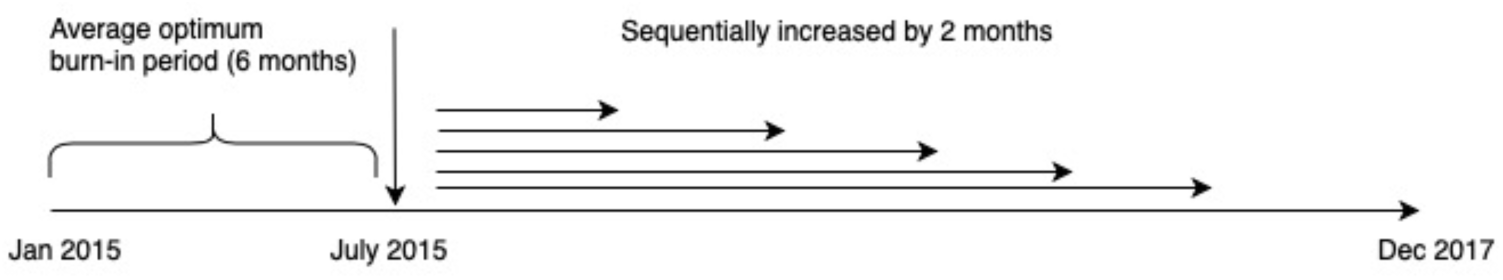
Modified approach used to assess the extent of prediction in aggressive outcomes by varying the testing dataset length.

Ranks predicted by the training dataset were then compared to the observed outcomes in each testing dataset. Percentage accuracy was plotted as a function of testing dataset length, giving us an insight into the rate of decay in each method’s performance.

### e. Elo-rating approach using optimized k

Rather than arbitrarily attribute k values to aggressive categories, we used an unsupervised way to assign them. The idea here is to find the values to fit each of the aggressive categories in our dataset. As we are seeking to find the optimised k values that fit best our data, we performed an optimization. We used the DEoptim package (Mullen et al. 2011). The DEoptim function searches for the global optimum of the objective function (fn) between lower and upper bounds on each parameter to be optimized. It is important to emphasize that the result of DEoptim is a random variable, i.e., different results may be obtained when the algorithm is run repeatedly with the same settings. In our case, the function fn with the highest percentage of accurately predicted outcomes was kept, along with the optimized parameters that corresponded to our four different categories of threat. We assigned to these parameters the lower bound of 0 and the upper bound of 500. Once the optimal parameter values had been extracted, individual ranks were calculated with the modified Elo-rating function provided by Newton-Fisher (2017).

The use of this optimization led us to modify our training/testing approach into a training/validation/testing one. Specifically, we divided the original 80% training dataset in two datasets, commonly called training and validation. The training dataset (i.e., the first 80%) was used to attribute k values, leading to the calculation of individual ranks based on these values (Figure 5). The remaining 20%, the validation dataset, allowed us to see how well these ranks did in predicting the aggressive outcomes. Depending on the percentage of accurately predicted outcomes, k values were updated accordingly in the training dataset. Once the optimised k values were obtained, they were used to calculate the ranks from the original 80% training dataset. The testing dataset then allowed us to test the efficiency of the calculated ranks in predicting future aggressive outcomes.

**Figure 5:**
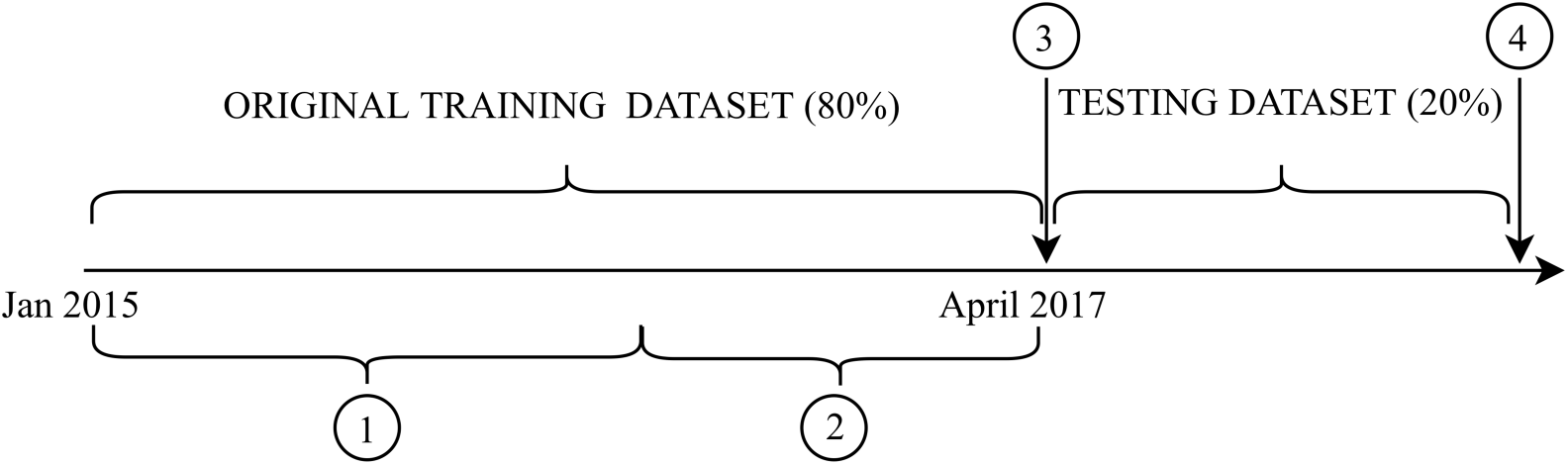
Modified approach used to improve the performance of the modified Elo-rating (1) Training dataset (80%) used to (i) attribute k values to the different types of aggression (ii) calculate individual ranks. (2) Validation dataset (20%) used to aggressive outcomes and allow the loop to modify the k values in order to improve the prediction. (3) Once the optimized k values obtained, final ranks are computed using the original training dataset. (4) Percentage of accrately predicted outcomes is extracted after testing the ranks on the testing dataset.

### Ethical note

All protocols were non-invasive and adhered to the laws and guidelines of South Africa and Canada. Procedures were approved by the University of Lethbridge Animal Welfare Committee (Protocols 0702 and 1505).

## RESULTS

### a. Which method performs best?

Our dendogram identifies the extent to which the methods provide similar estimates of rank order in our study group (Figure 6a). The height of the fission on the vertical axis indicates the similarity of ranks between two methods. The higher the height of fusion occurs, the less similar the methods are in terms of their outputs. With this in mind, the output from the original Elo-rating appears to be most different from the others, followed by the modified Elo-rating and then the I&SI method. The green cluster comprising the David’s scores method is the most similar in its outputs followed by the P&C method and then by the BI and randomized Elo-rating cluster.

**Figure 6:**
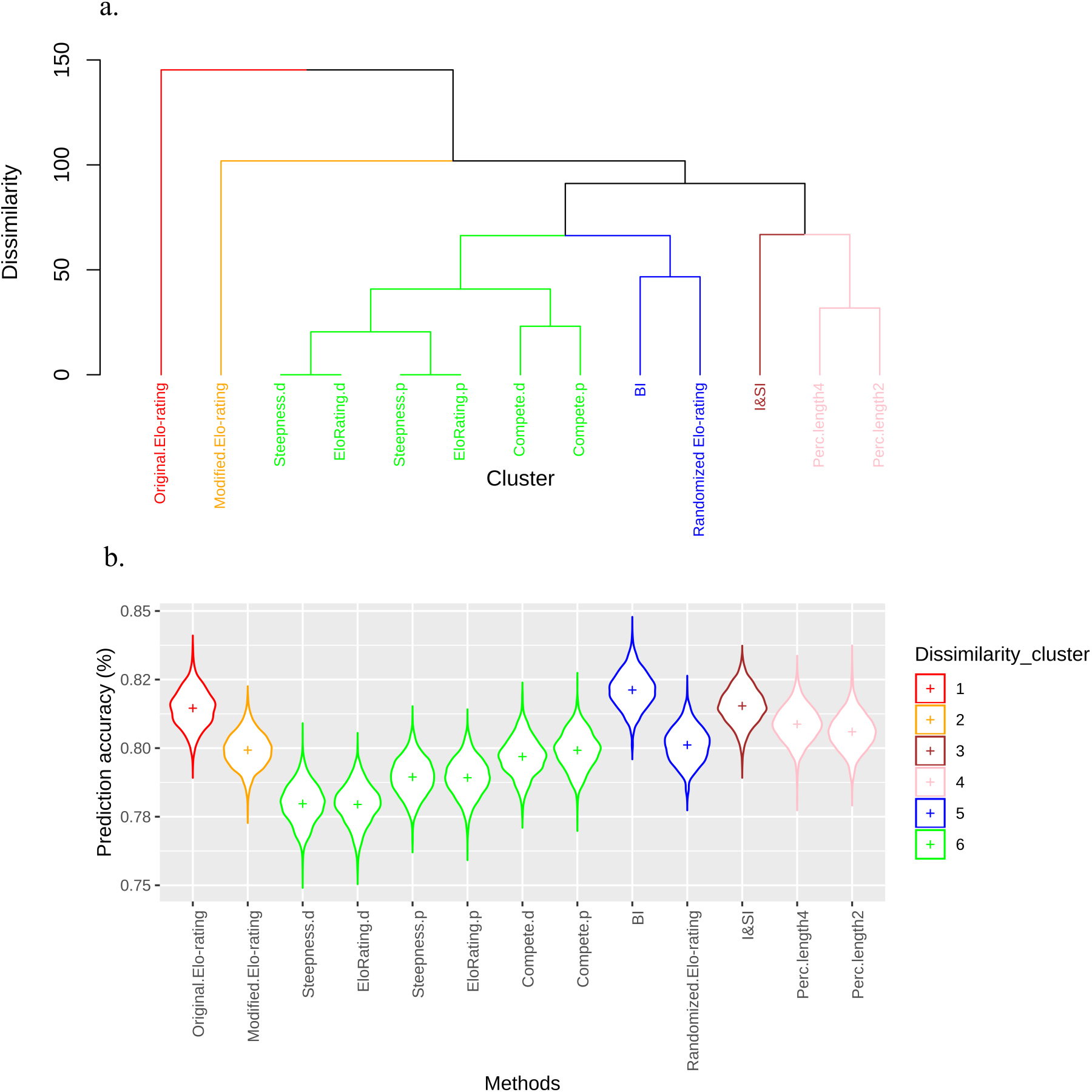
(a) How similar are the rank orders produced by each method? (b) How similar are the methods in their performance? Distribution of the percentage of accurately predicted outcomes across the methods used. Red, orange and blue clusters represent the interaction-based approaches while pink, brown and green ones are matrix-based methods. Green corresponds to the David’s score variants.

The violin plot, which shows the percentage of accurately predicted outcomes in the testing dataset, presents interesting results when combined with the dendogram (Figure 6b). While the dendogram shows how similar the methods are in their outputs (rank order), the violin plots show the variance in the percentage of accurate predictions produced by each method, i.e., they give us a sense of the “uncertainty” in the rank outputs produced. Looking at the blue cluster (BI and the randomized Elo-rating), we can see that both methods produce similar outputs (Fig 6a) and, yet, differ in their performance: the BI approach has a higher percentage of accurate predictions than the modified Elo-rating (Fig. 6b). Interestingly, the blue cluster of the BI/randomized Elo-rating and the modified Elo-rating (orange cluster) differ in their outputs (Fig 6a), but the randomized Elo-rating method’s performance is more similar to the modified one than the BI method (Fig 6b).

The percentages of accurately predicted outcomes for each method are given in Table 2. These indicate that all methods performed well in inferring reliable ranks (ranging from 77.5% to 82.9%). The I&SI method provided the best fit to our data, accurately predicting 82.9% of aggressive outcomes, with the BI approach producing an almost identical value of 82.6%. The David’s score obtained from the three different packages, and via the two different functions (Dij and Pij), were the lowest performing with the percentage of accurate prediction ranging from 77.5% to 79.9%, as well as the modified Elo-rating with a percentage of predicted outcomes of 79.9. The David’s scores from the “EloRating” package and the ones from the “steepness” package (Pij and Dij function) gave the exact same percentage outcomes. Compared to Dij function, the Pij predicted a higher number of accurate outcomes across all three packages used. Moreover, the “compete” package with Pij has the same efficiency as the k modified Elo-rating approach (79.91%). The I&SI method apart, the interaction-based approaches outperformed the matrix-based ones. Taken together, these results show that, despite these methods differing in their approach and the nature of their outputs, they all show a high level of efficiency in predicting future outcomes of aggressive interactions in our dataset.

**Table 2:** Percentage in aggressive outcome prediction over an 8-month testing dataset for our vervet monkeys’ troop (RBM).

| Method | Package | Option | % prediction<br>RBM |
| --- | --- | --- | --- |
| I&SI | Compete | isi13 | 81.9 |
| David's scores | Compete | Dij | 79.6 |
|  |  | Pij | 79.9 |
|  | Steepness | Dij | 77.5 |
|  |  | Pij | 78.7 |
|  | EloRating | Dij | 77.5 |
|  |  | Pij | 78.7 |
| Percolance &<br>Conductance | Perc | maxLength4 | 81.1 |
|  |  | maxLength2 | 80.7 |
| Elo-rating | EloRating | default | 81.8 |
|  | Newton-Fisher's code | K variation | 79.9 |
| Bayesian<br>Inference | Goffe's code | Rstan | 82.6 |
| Randomized elo-rating | aniDom | n.rands=1000 | 80.9 |

### b. Burn-in period: are some methods more efficient than others?

We sought to find a balance between using too few and too many data in our training dataset. We expected to see that, as the length of our training dataset increased, so the percentage of accurately predicted aggressive outcomes should also increase. However, as the training dataset begins to accumulate older aggressive interactions, we expected the accuracy of the percentage of predicted outcome to decrease. This assumption was borne out by our results to a certain extent (Figures 7 and 8). For the most part, however, accuracy of predicted outcomes was not greatly affected by the length of the training dataset.

**Figure 7:**
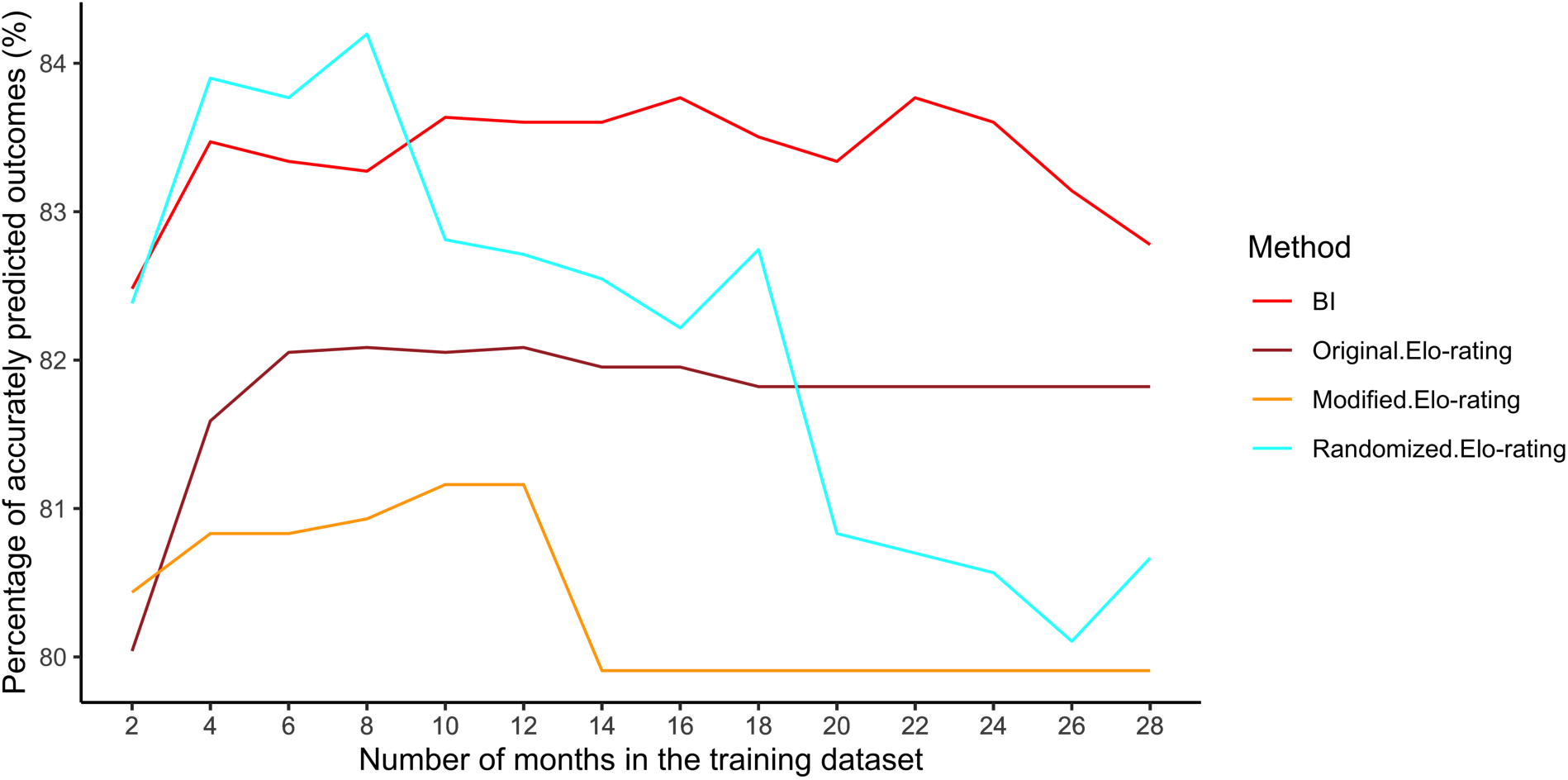
Variation of the percentage of outcome prediction with the dynamic approaches in function the number of months included in the training dataset.

**Figure 8:**
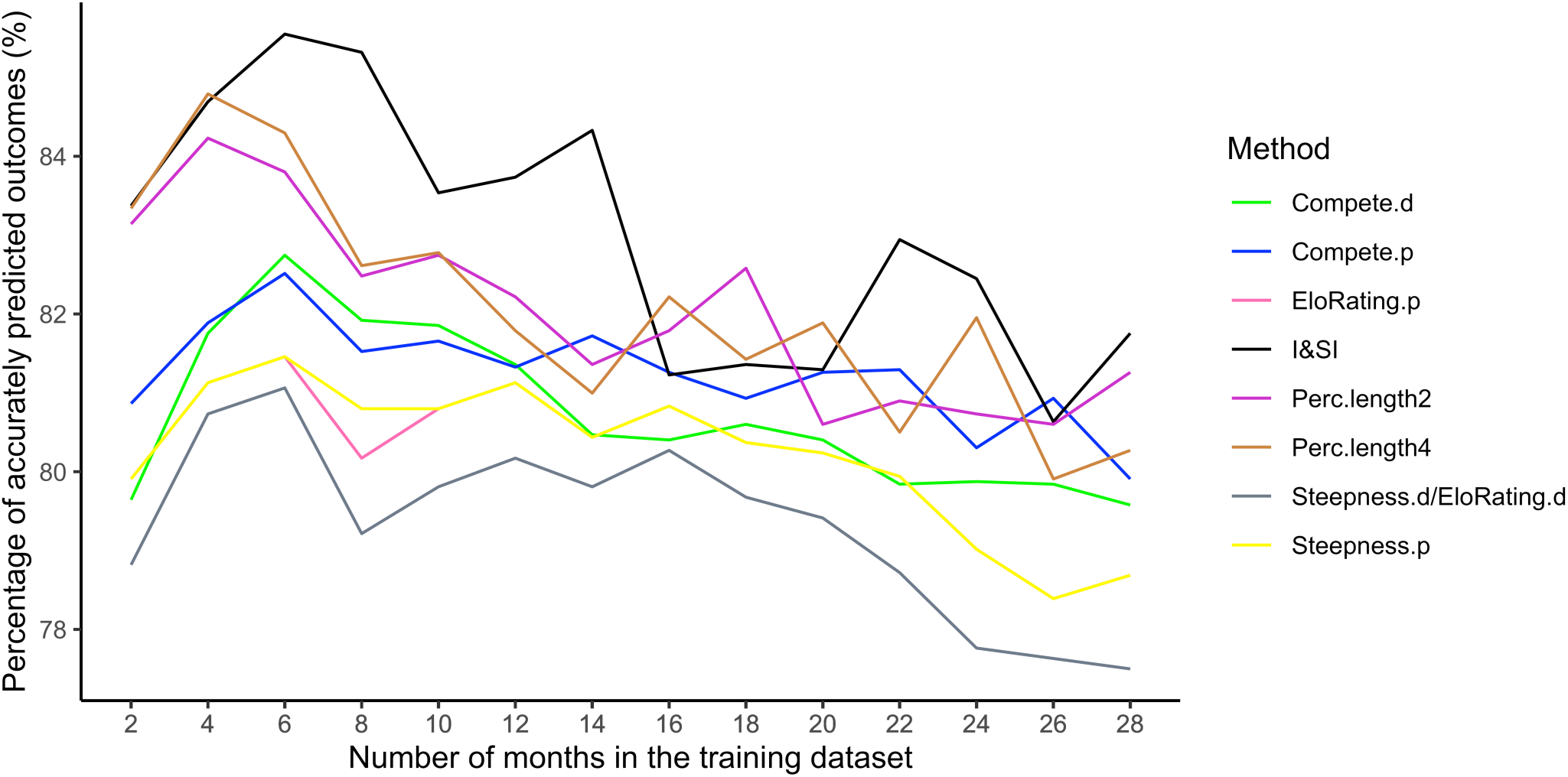
Variation of the percentage of outcome prediction with the static approaches in function the number of months included in the training dataset.

For ease of presentation and interpretation, we chose to look at the dynamic and static approaches separately. Figure 7 presents the dynamic approaches and shows that the length of the training dataset does not have any impact on three of the four methods: the original Elo-rating, the BI and the modified Elo-rating. In fact, these three methods perform very well regardless of the length of the training dataset. There is some evidence to suggest, however, that the BI approach and original Elo-rating method produce slightly better predictions once the training dataset has exceeded four months. As for the randomized Elo-rating method, this shows a more acute sensitivity to the amount of data in the training set. A first peak in accurate prediction appears at training set lengths between 4 and 8 months. Past 8 months, the accuracy in outcome prediction decreases as the training dataset length increases.

The static approaches (Figure 8) display much more variation in the percentage of accurately predicted outcomes depending on the length of the training dataset. Despite this sensitivity, a maximum in prediction accuracy can be found for each of these methods. On average, these peaks occur at 6 months, which we suggest represents the optimal length of burn-in period needed to accurately predict future outcomes in the data. As the number of months in the training dataset increases, we also see more of a decay in accurate prediction compared to the more dynamic methods (as one would expect). Moreover, the I&SI method displays a higher percentage of accurately predicted outcomes compared to the dynamic methods when the training dataset spans the period of 2 to 12 months.

### c. Ranks stability: increasing the testing dataset

To construct our training dataset for this analysis, the start date was determined by the average optimal burn-in period obtained from the previous analysis. With the exception of P&C, all methods from the static approaches had an optimal percentage of accurate prediction with a 6-month burn-in period. The dynamic approaches reached saturation sooner, between 2 and 4 months, the randomized Elo-rating being the exception with an 8-month burn-in period. We therefore used a 6-month burn-in period as this represents the best compromise in terms of enabling comparison across all methods. Shortening the burn-in period in this manner this manner gave us a larger testing dataset of 30 months in total (2.5 years).

The percentage of accurate predictions for each testing dataset length is plotted in Figure 9a-c. All methods show the same pattern. First, a decline in outcome predictability occurs at 4 months, which is then followed by a peak in prediction accuracy corresponding to a testing dataset of 8-10 months in length (Fig.9a). Past 10 months, the accuracy of predicted outcomes shows a constant and slow decay. These fluctuations are more or less marked depending on the method used. This pattern (i.e., where the curve’s fluctuation is more marked) is picked up by all dynamic approaches, while some of the static ones (I&SI and P&C) display the same pattern in fluctuations for the first 10 months, but this is much less pronounced. The I&SI, as well as the P&C approach, stand out as the methods producing a set of ranks that will lead to the highest percentage of accurately predict outcomes over the whole testing dataset’s length (Figure 9.a), followed by the BI approach that presents an overall slower decay in prediction. The rest of the methods are clustered with a lower percentage of accurately predicted outcomes throughout the testing dataset’s length.

**Figure 9:**
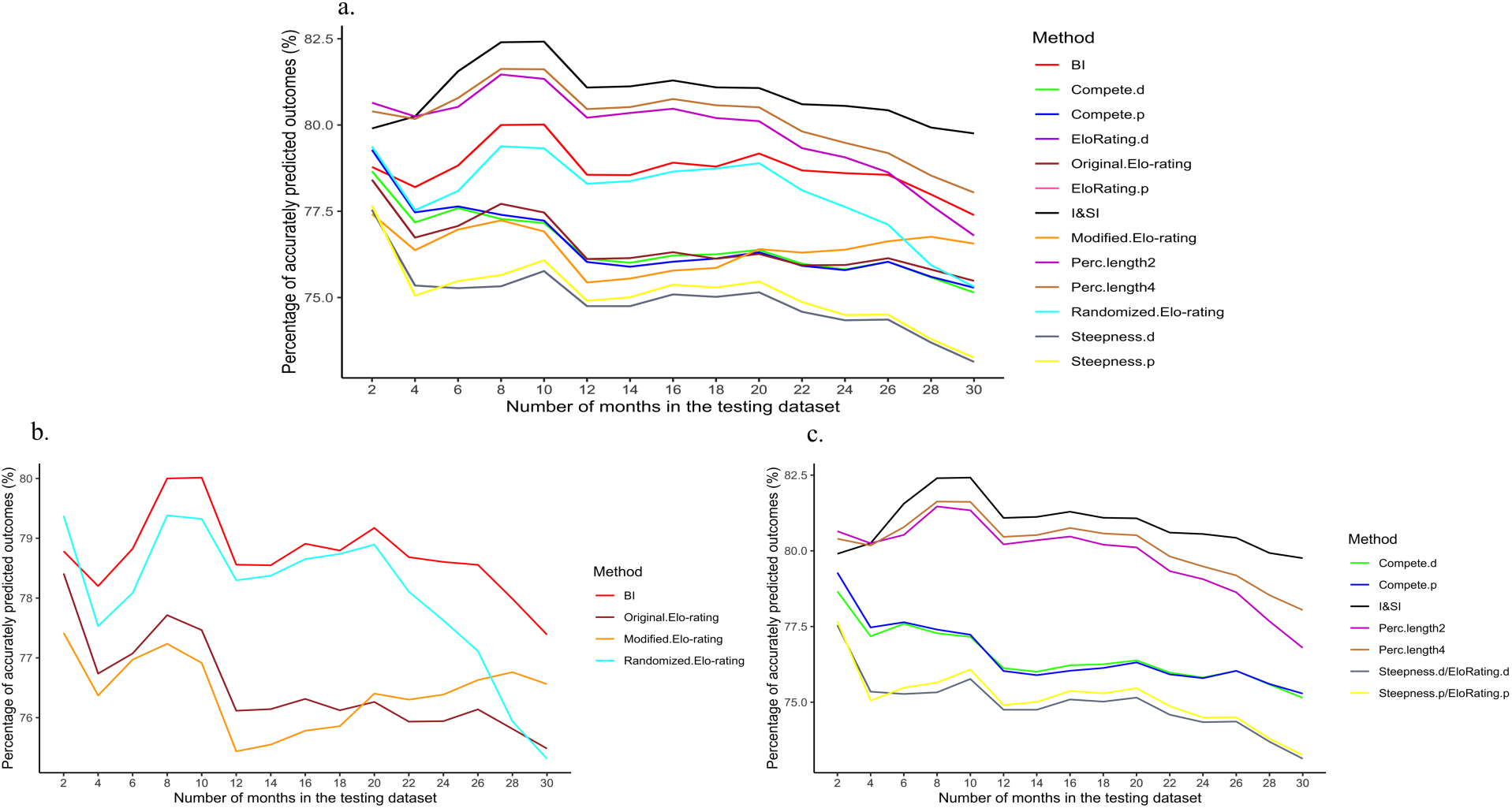
Variation of the percentage of accurate predictions as a function of testing dataset length across (a) all methods, (b) dynamic approaches only, (c) static approaches only.

In order to examine these patterns in more detail, we separated the dynamic and static approaches (Fig. 9 b and c). With respect to dynamic approaches, as mentioned above, the BI and the original Elo-rating display the general pattern described above. The randomized and the modified Elo-rating show lower performance overall but, following the peak in prediction accuracy at 10 months, the accuracy of the randomized Elo-rating then remains constant, while the modified Elo-rating produces an increasing percentage of accurately predicted outcomes as the length of the testing dataset increases. With regard to the static approaches, both the I&SI and P&C methods display the same pattern as the BI and the original Elo-rating. The David’s score shows a smoother curve, with a constant decrease in accuracy once past a testing dataset of 2 months. The “compete” package appears to perform better than the “steepness” and “EloRating” packages, which produce equivalent curves. Finally, the I&SI and P&C methods display a higher percentage of accurately predicted outcomes compared to the dynamic methods throughout the whole testing dataset length, except for P&C with maxLength 2 past 26-months of the testing dataset.

### d. A further modification to the modified Elo-rating

An optimization was performed on the last 20% of the original training dataset in order to obtain the best-maximised k values leading to the calculation of accurate ranks. The values attributed to the aggression categories are represented in Table 3.

**Table 3:** k values attributed to the four types of aggressive categories for the two different methods

|  | Modified Elo-rating | Alternative k values |
| --- | --- | --- |
| Physical aggression | 300 | 47.8 |
| Active aggression | 250 | 33.4 |
| Stationary aggression | 225 | 43.1 |
| Non-aggressive | 200 | 31.4 |

With these new k values, 84.3% of accurately predicted outcomes were obtained over the 8-month testing dataset. This alternative approach thus offers the best performance among all the tested methods. The use of the optimisation enabled us to increase its performance by nearly 5%.

## DISCUSSION

The short answer to our original question is that there is no best method per se. Rather, all methods performed well. Given this, it seems tempting to conclude that it is possible to simply flip a coin or otherwise choose a method randomly. This would be the wrong conclusion, however, as our analyses forced us to the realisation that the best method will always be a dataset-specific phenomenon. The true value of our analyses lies in the demonstration that a training-testing procedure provides researchers with the tool they need to guide their choice of method. In other words, although we have failed to identify *the* best method, we have generated a means by which researchers can find the best method for *them*.

More specifically, the training-testing procedure we used here not only enables informative comparisons to be made between different methods, but also gives researchers important insights into the length of the burn-in period needed to produce accurate rank predictions, as well as the future duration over which a given set of ranks can be used in analysis. In this way, researchers can determine which will be the most appropriate method, given sampling effort, the frequency of agonistic interactions, the steepness of the hierarchy, and the nature of the research question being asked.

The use of this procedure also allows researchers to probe the social dynamics of their study species in more depth. For example, when comparing ranks and outcomes, our assumption was that any mismatches represented a rank being wrongly attributed. However, it would be interesting (and possible) to look at the outcomes of aggressive encounters that do not match the ranks assigned to each participant, when both were extracted on the same day. This would give us a better idea of the true degree of outcome unpredictability, allowing us to assess whether uncertainty in rank assignment is due to the nature of the aggressive interaction itself or whether it reflects something about the context in which it takes place. Indeed, in cases of high outcome predictability and rank stability (i.e., where rank challenges would seem unprofitable), it may be the initiation and/or escalation of aggressive interactions that is unpredictable and linked to the broader ecological and social context. The procedure can also be used to assess the rate at which immature animals integrate into the adult hierarchy. In other words, the training/testing approach enables researchers to investigate dynamic pattern of hierarchy formation and maintenance, and is not simply a means of determining the most appropriate ranking method. Indeed, it can also be used to investigate temporal aspects of the aggressive outcome prediction at a dyadic level, rather than the whole group, and could thus offer a better understanding of the detailed social dynamics of a given study species. Another way to look at social dynamics is offered by Strauss & Holekamp (2019) who propose the concept of a longitudinal hierarchy to estimate the dynamics (Δ) of dominance hierarchies.

More generally, our findings demonstrate how researchers can use the training-testing procedure to choose the most appropriate method for their needs, given the amount of data they have at their disposal, and the nature of their study question. We acknowledge, however, that the use of empirical data does not allow us to distinguish between the two sources of error that could explain the methods’ performance: (i) inadequacies of the method, and (ii) real biological change. Thus our findings on how different methods compare are only valid with respect to our data and cannot be assumed to apply to other datasets. Within our dataset, we feel it is safe to assume that variation in a method’s performance compared to others reflects something about the method itself, given that we tested all methods on the same training/testing datasets such that any biological changes within the dataset should have be detected by all methods equally.

This is where the usefulness of simulated data comes into play as this allows one to tease apart these two sources of error more effectively, as well as gaining some more general insights into each method (Sánchez-Tójar, Schroeder & Farine, 2018). For example, Goffe, Fischer & Sennhenn-Reulen’s (2018) advocate running simulation approaches to test how well Elo-ratings reflect the (simulated) true structure and how sensitive the rating is to true changes in the hierarchy. Nevertheless, we believe the training-testing procedure is valuable empirically for those interested in determining how much the methods vary when run on the exact same dataset, and why this might be. It is also the case that, although simulated data provide insight into how well a given method is able to recover the true hierarchy, simulated data are also much tidier and “well behaved” compared to actual field data, so one should still be cautious about proclaiming any one method better than the others, as it may be sensitive to qualities of empirical data that do not occur in simulations.

With respect to our specific dataset, our main finding is that all methods tested performed well, despite some differences in inferred rank order. With respect to the required burn-in period, again, all methods appear to be efficient in inferring reliable ranks from the very start (i.e., with little to no burn-in period), but all showed improvement as the burn-in period increased. In general, we found that dynamic methods were less sensitive to the amount of data present in the training dataset compared to the static approaches. As a result, dynamic approaches show a constant efficiency regardless the length of the training dataset compared to the static approaches. This is not unexpected given that methods dynamic approaches track rank variations continuously and update the ranks after each interaction. We should also highlight that, when the training dataset did not exceed 12 months, it was the static I&SI and P&C methods that produced the highest percentage of accurately predicted outcomes.

When we adopted a common 6-month burn-in period, all methods could predict aggressive outcomes accurately for the subsequent 10 months. Past this 10-month threshold, however, accuracy in prediction decreased as the testing dataset increased in length, but the decay was rather shallow, and there was a high predictability in aggressive outcomes across time, indicating overall rank stability during this period. This is not to say, however, that rank predictability did not fluctuate as the length of the training period increased, with the degree of fluctuation was dependent on the length of the method used. This suggests that real rank shifts were occurring in the study group during particular periods, and that the training-testing procedure provides a way to home in on such periods of instability. The combination of high overall predictability in aggressive outcomes with some temporal fluctuation suggests that, in our study species, rank order shows a form of “regression to the mean”. That is, individuals may experience very mild shifts in relative rank position up or down the hierarchy across time, but nevertheless occupy more or less the same “absolute” position in the hierarchy. This, in turn, suggests that rank changes may reflect the ecological and demographic contexts in which they occur, rather than pointing to genuine changes in inherent power. In line with this interpretation, the dynamic methods produce more fluctuations than the static methods: the former are likely to catch small shifts in rank position as they constantly update, whereas the static methods, which combine and cluster interactions across time, are more likely to produce a rank order that fits the overall social dynamic.

It is also important to remember that we converted the output from each method to ordinal ranks to facilitate comparison, and differences between (the various kinds of) Elo-ratings can be absolutely very small. Such differences would lead to a shift in calculated rank order and hence to fluctuation in prediction accuracy, but, in reality, may have no discernable impact on an individual’s social standing in the group. That is, such fluctuation may, in effect, represent noise around a stable rank order. Even though Strauss & Holekamp (2019) recommend the use of ordinal rankings for identifying hierarchy dynamics, this point thus serves to highlight a limitation of our study in that we looked only at rank order and did not consider the magnitude of rank differences nor any uncertainty around rank calculations. These components may very well matter, especially in species where a linear rank order may not be representative of the social hierarchy. This point also serves to highlight the true advantage of methods like the BI, randomized Elo-rating and the P&C approach, which enable researchers to look at the uncertainty around ranks, and thus gain a better understanding of the social hierarchy.

Our training/testing approach procedure also allowed us greater insight into the working of certain methods. For example, the randomized Elo-rating approach was developed to enhance performance in societies where ranks remain fairly stable across time (Sánchez-Tójar, Schroeder & Farine 2018) —a situation that, as noted above, pertains in our study group. Under such conditions, the sequence in which interactions occur should not affect the inferred ranks. Our results help validate this assumption. When the training data set was short (i.e., 6 months), the sequence of interactions clearly did not matter, and the method gave good results. As the training dataset increased up to 28 months, however, there was a decline in accuracy, which no doubt reflects the fact that, over this period, demographic change was inevitable and the order of interactions may well have begun to exert an influence on the structure of the hierarchy. Given this outcome with our data (and assuming this holds true across other datasets), this suggests that the randomised Elo-rating method will also prove useful in determining when and where interaction order matters in a given dataset. Indeed, Sánchez-Tójar, Schroeder & Farine (2018) suggest that researchers can control for factors such as changing ranks or winner-loser effects by controlling which parts of the data are randomised. For example, researchers interested in tracking how individual ranks change across days could increase the accuracy of the daily rank estimates by randomizing observations within each day.

Our own modification to the Elo-rating also provided some insight into the method’s workings. We produced an increase of nearly 5% in accurately predicted outcomes compared to Newton-Fisher’s (2017) modified Elo-rating. Newton-Fisher’s (2017) argument was that allowing k to vary would improve accuracy as the specific nature of aggression could affect the outcome and impact of aggressive encounters. Our results did not support this, but we reasoned that this was most likely due to the fact that our values of k were assigned arbitrarily. Switching to a data-driven approach to assign k-values, via the training/validation/testing procedure, resulted in a marked improvement in performance. Thus, the type of aggression shown does, in fact, matter to predictive accuracy, but arbitrarily assigning values to aggression types may fail to represent its importance correctly. It is also worth noting that we did not investigate the extent to which k-values, and hence ranks, were affected by the relative length of the training and testing periods used in the optimisation procedure (i.e., we used only a 80:20 split), nor did we consider any potential variation in values of k through time (e.g., due to seasonal effects). More work is needed to understand better the impact of relative lengths of training and testing periods, as well as the stability of k values through time.

In conclusion, we suggest that the insights into group dynamics provided by a training-testing procedure, combined with the ability to determine which ranking method is best suited to a given dataset, far outweigh the time costs associated with running the procedure. A training-testing approach gives researchers the power to tailor the selection of a dominance-ranking method to the particular nature of the dataset they are using, allowing them to take into account sampling effort, frequency of agonistic interactions and the steepness of the hierarchy, as well as the particular kinds of analyses they wish to conduct. In addition, comparison of methods and the dynamic patterns they reveal further enable researchers to home in on regions of their dataset that will permit analyses into how and why rank shifts occur and discover the underlying causes of both rank stability and unpredictability across time.

## Acknowledgments

We thank Mark and Sarah Tompkins for the permission to work at Samara, Kitty and Richard Viljoen for their continued logistic support. We thank Damien Farine and Alfredo Sánchez-Tójar for helpful discussions as well as for pointing out an error in the rank’s computation with the randomized Elo-rating method. We are also very grateful to the many research assistants who contributed to the database, as well as our amazing monkeys (and George), without which only simulated data would be available.

The research was supported by NSERC Discovery (Canada) grants to LB and SPH and by the Canada Research Chairs program to LB, TRB and CV.

## Appendix

The following sections explain in more detail the choices made regarding the analyses for each method.

### The I&SI method

It is recommended to find the matrix with the lowest SI associated with a certain number of iterations (nTries). To do so, we performed an optimization to find the nTries that best fit our data. We used the DEoptim package (Mullen et al. 2011). The DEoptim function searches for the global optimum of the objective function (fn) between lower and upper bounds on each parameter to be optimized. It is important to emphasize that the result of DEoptim is a random variable, i.e., different results may be obtained when the algorithm is run repeatedly with the same settings. In our case, the function fn with the highest percentage of accurately predicted outcomes was kept, along with the optimized parameter corresponding to the number of iterations (nTries). We assigned to this parameter the lower bound of 50 and the upper bound of 1000. Once the optimal parameter value had been extracted, individual ranks were calculated using the latest function version “isi13” from the R package “compete”.

The use of this optimization led us to modify our training/testing approach into a training/validation/testing one. Specifically, we divided the original 80% training dataset in two datasets, commonly called training and validation. The training dataset (i.e., the first 80%) was used to attribute the nTries value, leading to the calculation of individual ranks based on this value. The remaining 20%, the validation dataset, allowed us to see how well these ranks did in predicting the aggressive outcomes. Depending on the percentage of accurately predicted outcomes, the nTries value was updated accordingly in the training dataset. Once the optimised nTries value was obtained, it was used to calculate the ranks from the entire, original 80% training dataset. The testing dataset then allowed us to test the efficiency of the calculated ranks in predicting future aggressive outcomes. Here nTries= 449 was chosen.

### David’s Score

This approach proposes two alternative indices to compute David’s scores: Pij and Dij. Pij represents the winning proportion of individual i against j, which leads to a matrix of observed win proportions as an output. For the Dij index, a matrix is obtained where the observed proportion of wins (Pij) is corrected for the chance occurrence of this observed outcome. Balasubramaniam et al. (2013) argued that Pij might be a better choice for species with high levels of directional asymmetry (i.e., despotic species), whereas Dij may be a better choice for species with low levels of directional asymmetry (i.e., tolerant species). We compared both these indices. Furthermore, the David’s score method can be calculated with the aid of three different R packages. The decision was made to include them all in the analysis, producing two calculated scores per package. This allowed us to assess whether the calculated ranks were the same across all packages and if not, which package led to the highest percentage of accurate predictions.

### Percolation and Conductance

The parameter maxLength helps find all indirect pathways of a particular length and then update the conflict matrix. Examining information gained through indirect pathways provides information that can be used to decide on the appropriate maxLength for a dataset. To assist with this decision, the Perc package offers a transitivity function as a way to estimate an alpha value, which is used to weight the information from the indirect pathways to give an indication of the extent to which we can trust information from indirect pathways. Greater transitivity is associated with assigning higher weight to information from indirect pathways (Fushing, McAssey & McCowan 2011). We tested MaxLengths of 2 and 4.

### Elo-rating method

In the original method, k is held constant and all individuals receive the same elo-rating at the initiation of the burn-in period. Elo-ratings were calculated with 1000 as the initial value and k set to 100 (Neumann et al. 2011).

*1. <u>Derivations of the Elo-rating methods</u>*

### Modified Elo-rating (k variation)

In this method, each dominance interaction is classified according to the most intense level of aggression displayed by the winner (Newton-Fisher 2017). This being required, we excluded behaviours recorded as “unknown” from the analyses and distinguished among non-aggressive, stationary, active and physical threats. This classification is based on our inter-troop encounter protocol (Barrett, L. and Henzi, S.P., unpublished data). Non-aggressive behaviours included supplants (i.e., where the aggressor takes the victim’s place) and displacements (i.e., when one animal submissively moved away when approached within 10 meters). Any aggressive behaviour that did not include a forward movement was considered a stationary threat, such as lunge, facial and vocal threats. Active aggression involved ongoing forward movement (i.e., chase or charge) but where no physical contact was made with the target of aggression. Physical aggression was scored in instances where body contact was made (e.g., a bite or slap). We assigned a different K value to each of these categories, using the default value of 200 (Neumann et al. 2011) for the most commonly observed form of aggression (i.e., the non-aggressive interactions of displace and supplant) and scaling up in multiples of 25 to distinguish varying intensities. This led to the creation of a modified training data set and its detailed composition is given in Table A1.

**Table A1:** Modified training data set for the modified Elo-rating method (Newton-Fisher 2017).

|  | Modified training data set |  |  |  |  |
| --- | --- | --- | --- | --- | --- |
| Aggression Number | Total | Physical | Active | Stationary | Non-aggressive |
| RBM Troop | 8083 | 886 | 1644 | 1625 | 3934 |
| K value | - | 300 | 250 | 225 | 200 |

### Bayesian inference (BI) approach

We implemented this method Goffe, Fischer & Sennhenn-Reulen’s (2018) code (with no additional analytical choices or justification required).

### Randomized Elo-rating

By randomising the order of observed interactions, this approach allows the creation of K replicated datasets, where K corresponds to the number of randomisations performed. In this study we were only interested in obtaining the final scores (return.as.ranks =TRUE). The function returns a NxK matrix that gives the final scores for each individual (rows) after each randomisation of the order. In order to use the information contained in all the iterations, we extracted the mean ranks for each individual.

We followed the same procedure as Sánchez-Tójar, Schroeder & Farine (2018) and randomized the order in which interactions occurred 1000 times.

## Notes

#### Summary of Updates

The paper was re-arranged: an appendix was added.

